# Passive sampling of environmental DNA in aquatic environments using 3D-printed hydroxyapatite samplers

**DOI:** 10.1101/2021.05.12.443744

**Authors:** Héloïse Verdier, Lara Konecny, Christophe Marquette, Helen Reveron, Solène Tadier, Laurent Grémillard, Amélie Barthès, Thibault Datry, Agnès Bouchez, Tristan Lefébure

**Author notes:** Corresponding authors: Héloïse Verdier, Univ Lyon, Université Claude Bernard Lyon 1, CNRS, ENTPE, UMR 5023 LEHNA, Villeurbanne, France, Tristan Lefébure, Univ Lyon, Université Claude Bernard Lyon 1, CNRS, ENTPE, UMR 5023 LEHNA, Villeurbanne, France.

## Abstract

1. The study of environmental DNA released by aquatic organisms in their habitat offers a fast, non-invasive and sensitive approach to monitor their presence. Common eDNA sampling methods such as filtration and precipitation are time consuming, require human intervention and are not applicable to a wide range of habitats such as turbid waters and poorly-accessible environments. To circumvent these limitations, we propose to use the binding properties of minerals to create a passive eDNA sampler.
2. We have designed 3D-printed samplers made of hydroxyapatite (HAp samplers), a mineral known for its high binding affinity with DNA. The shape and the geometry of the samplers have been designed to facilitate their handling in laboratory and field. Here we describe and test the ability of HAp samplers to recover artificial DNA and eDNA.
3. We show that HAp samplers efficiently recover DNA and are effective even on small amounts of eDNA (<1 ng). However, we also observed large variations in the amount of DNA recovered even under controlled conditions.
4. By better understanding the physico-chemical interactions between DNA and the HAp sampler surface, one could improve the repeatability of the sampling process and provide an easy-to-use eDNA sampling tool for aquatic environments.

## 1 INTRODUCTION

At a time of unprecedented threats on freshwater biodiversity, it is crucial to develop rapid, accurate and minimally invasive tools to monitor aquatic ecosystems. About a decade ago, methods based on the sampling of environmental DNA (eDNA) were proposed as a revolutionary way to survey aquatic macro-organisms (Deiner et al., 2017). Macro-organisms release DNA in their environment through different processes (e.g. faeces, excretion, shedding cells, gametes) and this eDNA can take different forms (tissues, cells, organites, nucleo-proteic complexes, …). The direct sampling of eDNA coupled with molecular analysis methods such as NGS (Shokralla et al., 2012) or quantitative polymerase chain reaction (qPCR) (Langlois et al., 2020) allow the detection and identification of aquatic species while overcoming organism capture. Although eDNA offers many promising applications, several methodological challenges remain.

One of the most challenging aspects of eDNA-based approaches is the sampling method. eDNA is present in very small quantities and is heterogeneously distributed in aquatic environments (Goldberg et al., 2016). To maximise its recovery, sampling methods must be able to concentrate eDNA (Hinlo et al., 2017). Active filtration of a large volume of water is the most commonly-used method to recover eDNA in aquatic systems. However, filtration has significant methodological limitations. Firstly, it is a long and tedious process requiring human intervention, sometimes difficult to carry out in poorly-accessible habitats. Secondly, the clogging of the filters is a recurrent problem which reduces the volume of water that can be sampled (Williams, Huyvaert and Piaggio, 2017).

To limit clogging, filtration membranes with large porosity (greater than 0.45 μm) are often used. However, eDNA particles can be present in various forms (intra or extracellular), states (free or complexed with other particles) and sizes (from > 180 to < 0.2μm but most abundant between 0.2 and 10 μm) (Turner et al., 2014; Moushomi et al., 2019; Wilcox et al., 2015). As filtration is based on particle size sorting, the use of membranes with large porosity will overlook smaller DNA particles, even though they may be an important source of eDNA (Moushomi et al., 2019). Finally, given the complex dynamic of eDNA in aquatic environments (i.e. pulsed emission, transport, retention, degradation), one filtration sample will provide an instantaneous snapshot which is likely to be poorly integrative of the overall eDNA signals (Pilliod et al., 2013; Spear et al., 2015).

Passive eDNA sampling using natural substrates is a promising solution to overcome filtration challenges. Organisms such as marine sponges (Mariani et al., 2019), molluscs (Der Sarkissian et al., 2020) and biofilms (Rivera et al., 2021) can trap and accumulate eDNA particles in water. Minerals can also accumulate and protect DNA from enzymatic degradation (Alvarez et al., 1998; Levy-Booth et al., 2007). Indeed, a sample of sediment can contain more eDNA than a water sample (Turner, Uy and Everhart, 2015). Recently, Kirtane and colleagues (Kirtane, Atkinson and Sassoubre, 2020) have shown that montmorillonite and coal-based mineral powders can be used as passive eDNA samplers in aquatic environments. Thanks to good DNA capture and preservation rates (up to 200 μg genomic DNA / g) (Gardner and Gunsch, 2017), sediments and commercial mineral powders may very well be more integrative eDNA substrates than filtration methods. Yet, these substrates are difficult to handle and deploy in the environment, particularly in aquatic systems.

In this study, as an alternative to filtration, we developed proof of concept 3D-printed passive eDNA samplers. 3D printing allows control of the shape and composition of an object. The shape of the samplers were designed with optimised surface/volume-ratio and a shape easing handling in the field and in the lab. The samplers were made of pure hydroxyapatite (HAp), a calcium phosphate mineral naturally present in bones and known for its high binding affinity toward DNA (Okazaki et al., 2001; Brundin et al., 2013). Here we describe the development of hydroxyapatite samplers (HAp samplers) and test their ability to sample eDNA in fresh waters. Two prototypes of samplers will be presented: a first test-version, with which the concept and material will be tested, and a second version which shape and design have been optimised for eDNA sampling. Using controlled laboratory experiments, our objectives are to (i) quantify the HAp samplers DNA binding and release capacity, (ii) assess the range of DNA fragment size sampled, (iii) quantify the repeatability of DNA sampling across several cycles of use of the HAp samplers, and (iv) evaluate the samplers capacity to sample eDNA released by organisms in microcosm.

## 2 MATERIALS AND METHODS

### 2.1 3D-printed HAp samplers design

#### 2.1.1 Raw material and printing setup

A photopolymerizable organic resin (3D Mix, 3DCeram Company, HAP, Bonnac-la-Côte, France) containing 40-60 % (w/v) of hydroxyapatite powder (Ca_10_(PO_4_)_6_(OH)_2_, stoichiometric hydroxyapatite), a synthetic calcium phosphate with Ca/P atomic ratio of 1.67, was the raw material used to fabricate the samplers. The samplers were built from this hydroxyapatite-enriched resin using a 3D stereolithographic printer (CERAMAKER C900, 3DCeram Company, with 55mW laser power and 100μm layer thickness).

Two types of prototypes of HAp samplers were produced. The first prototype (P1) is a test version corresponding to 10 pieces cut out of a 3D-printed mesh (Fig. 1a). P1 prototypes have an exposed surface of 240 mm^2^ and a macroporosity of 500 μm in diameter. A second more elaborate prototype (P2) was then produced with optimized geometry and porosity, and printed in 25 copies (Fig. 1b). P2 has a total surface of 480 mm^2^ and a macroporosity of 400 μm in diameter.

**FIGURE 1:**
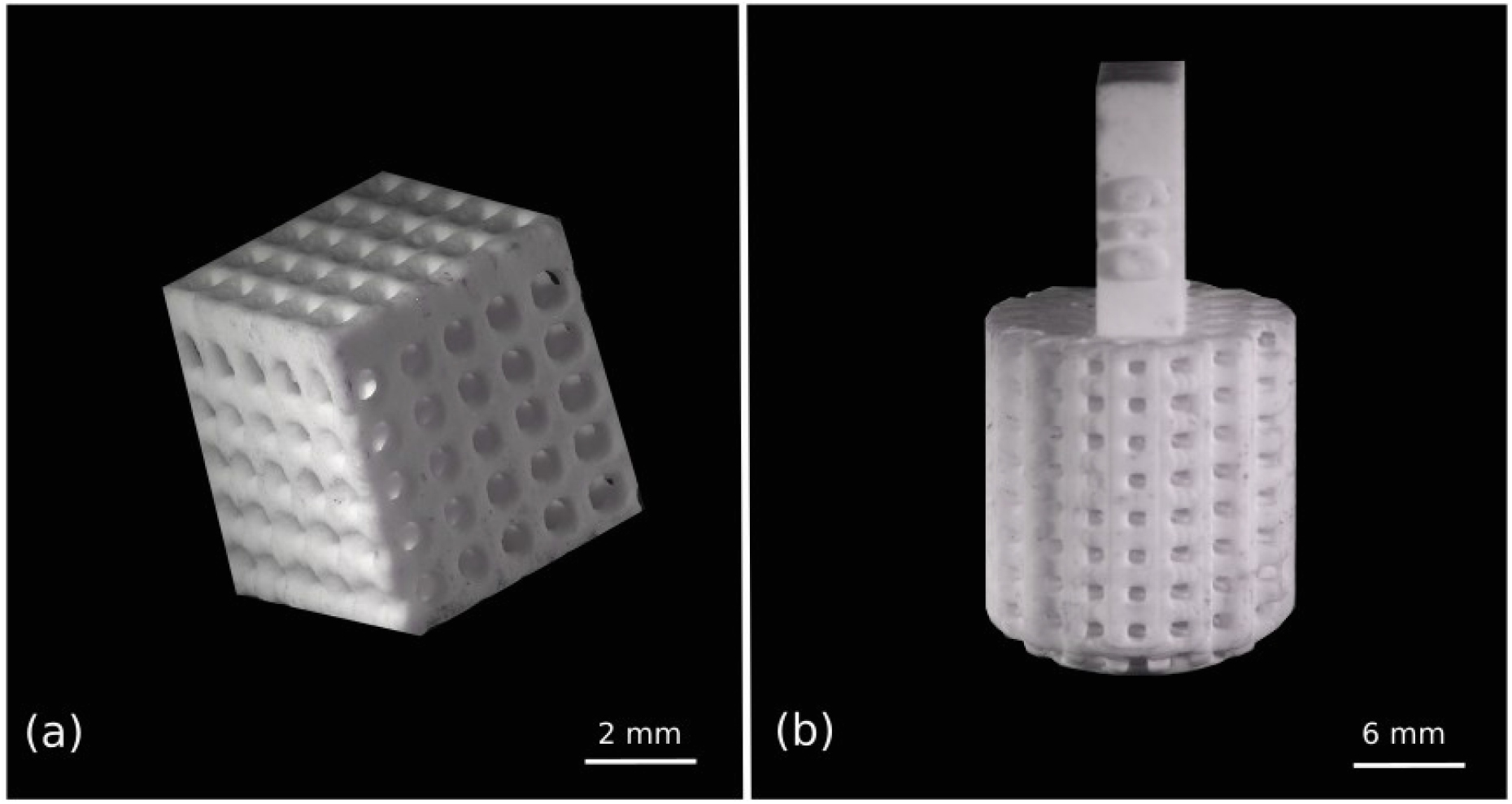
Images of 3D-printed hydroxyapatite samplers prototype P1 (a) and P2 (b) obtained with a confocal microscope (objective x0.5, LEICA Z16 APO, camera LEICA DMC5400).

#### 2.1.2 Debinding and sintering steps

Once printed, cleaned with a specific solvent (Ceracleaner, 3DCeram Company, Bonnac-la-Côte, France) and dried, the HAp samplers underwent debinding and sintering steps. Debinding aims at removing all organic components (in particular the organic resin) and was conducted in a conventional oven following the thermal cycle described in Table 1.

**TABLE 1:** Process parameters for debinding HAp samplers

| Step | Temperature (°C) | Heating rate<br>(°C/min) | Dwell (min) |
| --- | --- | --- | --- |
| 1 | 20-200 | 0,2 | 120 |
| 2 | 200-300 | 0,1 | 120 |
| 3 | 300-380 | 0,1 | 120 |
| 4 | 380-550 | 0,1 | 120 |
| 5 | 550-950 | 1 | 0 |
| 6 | 950-20 | 2 | - |

Sintering aims at consolidating the samplers by densifying them (creation of necks and reduction of the porosity between the individual ceramic particles) (Rahaman, 2017), and is achieved by a thermal treatment at higher temperature (1°C/min up to 1150°C, 60 min. at 1150°C, followed by a second step at 3°C/min up to 1250°C, 60 min at 1250°C, finally cooling to room temperature at 3°C/min). After sintering, no additional processing (i.e. finishing or polishing) was performed. The presence of pure HAp was confirmed by X-ray diffraction (XRD) performed on as-sintered samples.

### 2.2 Expected DNA recovery from HAp samplers

We used the term “DNA recovery” to define the quantity of DNA that binds and is released from the HAp samplers. We estimated the theoretical maximum DNA recovery (DNA_max_) based on the hypothesis that a single layer of DNA molecules would bind on the HAp surface of the samplers. According to equation 1, the number of DNA molecules that can bind to the surface is obtained by dividing the exposed surface (Se) of a sampler (P1 = 240 mm^2^, P2 = 480 mm^2^) by the surface of a DNA base pair (Sd = 6.46E-10 mm^2^). The surface of a DNA base pair was calculated according to Mandelkern et al (1981) (diameter = 2 nm, length = 3.4 nm). The number of DNA molecules per sampler is then divided by Avogadro’s constant (NA = 6.02214076 × 1023 mol - 1) to give the number of DNA moles per sampler. The number of moles of DNA is then divided by the molar mass of a DNA base pair (W = 650 daltons) to obtain the total mass of DNA that can bind to a sampler. Being smaller, P1 has a maximum theoretical recovery capacity of 400 ng of DNA per sampler, while P2 has a capacity of 800 ng.

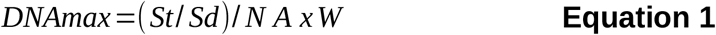

### 2.3 Protocol of DNA binding and release

The HAp sampler DNA binding and release protocol is composed of 5 steps. First, HAp samplers are decontaminated before each experiment by a thermal treatment in air at 550 ° C for 3 hours (Thermolyne model 30400 furnace), a procedure typically used to decontaminate glassware. Second, DNA is bound to the HAp samplers by immersing them in an aqueous solution (varying composition upon the present study) containing DNA. Third, samplers are transferred to Eppendorf tubes and centrifuged for 1 minute at 3000 rpm to dry them. Fourth, samplers are washed with 1 mL of sterile ultrapure water. Finally, DNA is released from the samplers by immersing them in 1 mL of 0.1 M phosphate buffer pH 8, vortexed for 30 seconds and incubated at room temperature for 1 hour.

**FIGURE 2:**
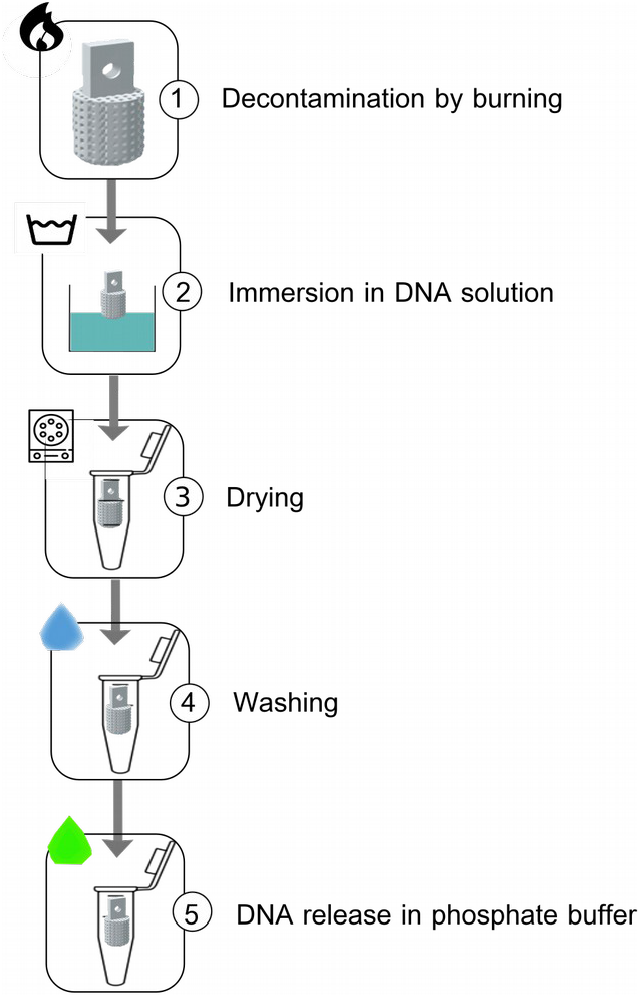
DNA binding and release protocol

### 2.4 DNA sampling experiments by HAp samplers

#### 2.4.1 Experiment 1: DNA binding and release

A DNA sampling experiment with concentrated DNA fragments of various sizes (i.e. a DNA size marker) was conducted to validate the binding and release protocol and to assess whether DNA fragments of different sizes have different binding efficiencies. After decontamination, one batch of HAp samplers (P1 and P2) was incubated in tubes (1 sampler/ tube) containing 2 mL of a solution of large DNA fragments at 5000 ng / mL (λ DNA / BstEII Digest, 117-8450 bp). A second batch of HAp samplers was incubated in tubes containing 2 mL of a solution of short DNA fragments at 2000 ng / mL (PCR 20 bp Low Ladder, 20-2000 bp). Both batches were incubated for 17 hours on a rotary shaker (IKA Roller 6 Digital, 40 rpm). Controls were tubes with 2 mL of solution and devoided of samplers. Residual DNA in the supernatants was quantified by taking 60 μl aliquots of the supernatant after 45 min and 17 h of incubation. After incubation, HAp samplers were removed from the DNA solutions using sterile clamps and DNA was released according to the protocol in section 2.3. All supernatants aliquots and released DNA solutions were stored at −20°C prior to analysis.

#### 2.4.2 Experiment 2: repeatability

A quantification of repeatability was conducted to test whether HAp samplers can be reused after several cycles of use. A cycle of use is defined here as a thermic treatment phase followed by a DNA binding and release phase. For this purpose, 5 prototypes 1 and 25 prototypes 2 HAp samplers were incubated in a concentrated solution of DNA size marker (λDNA/BstEII Digest 117-8450 pb) at a concentration of 2.8 μg/mL on a rotary shaker (Roller 10 Digital IKA) for 17 H. This experiment was carried out three times in a row (hereafter called experiments A, B and C) under strictly identical conditions, at room temperature (24°C ± 2°C) with decontamination through thermic treatment between each use. After incubation, HAp samplers were removed from the DNA solution with sterile clamp, washed and DNA was released with 1 mL of 0.1 M phosphate buffer pH 8 according to the protocol section 2.3 DNA samples were stored at −20°C prior to analysis.

#### 2.4.3 Experiment 3: microcosm experiment

*Asellus aquaticus*, a small freshwater isopod, was used as a target organism to test the capacity of the HAp samplers to collect eDNA. 40 organisms of *A. aquaticus* sampled from a natural pond (Lyon, France) in april 2019 were divided into 8 glass microcosms (5 individuals / microcosm) containing 500 mL of synthetic water (Peltier and Weber, 1985). Positive controls correspond to microcosms where we injected genomic DNA (final microcosm at 1 ng/mL) extracted from a pool of 10 *A. aquaticus*. Negative controls were of two types: control microcosms containing water without DNA and a sampler, and control samplers from which the DNA was released just after the decontamination step (i.e. without incubation in DNA solution). After 24 hours of *A. aquaticus* acclimatization, the two prototypes of HAp samplers were incubated in microcosms (1 sampler / microcosm) for 24 hours. All microcosms were placed in a cold room at 18°C, spaced 0.5 m apart and covered to limit the risk of contamination. The organisms were not fed during the experiment to reduce the amount of allochthonous DNA. After incubation, the HAp samplers were collected with sterile clamps and the DNA was released according to section 2.3 of the protocol. Released DNA was purified (Macherey-Nagel ™ NucleoSpin ™ gel and PCR cleaning kit) to avoid potential inhibition of the downstream qPCR by the phosphate buffer (see next section), following the manufacturer’s recommendations. Purified eDNA was stored at −20 ° C prior to analysis.

**FIGURE 3:**
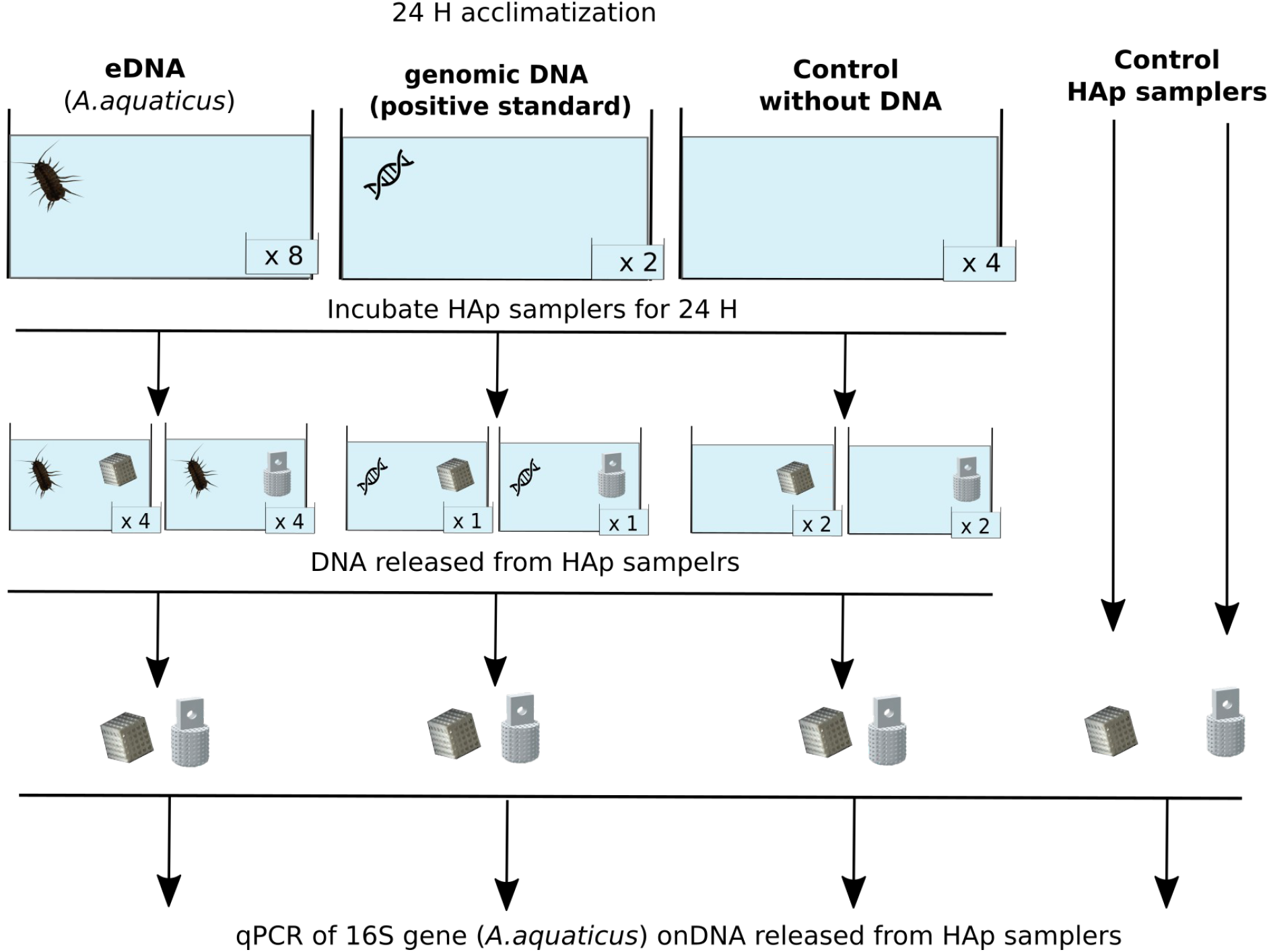
Experimental design testing HAp samplers efficacy to recover eDNA from *Asellus aquaticus* in microcosms.

### 2.5 DNA quantification and analysis

#### 2.5.1 Quantification of DNA size marker

In the first experiment, DNA binding and release by HAp samplers were evaluated by following the DNA concentration and fragment sizes in three compartments: (1) in the supernatant (i.e. residual DNA), (2) in the washing solution, and (3) in the releasing solution (Fig. 4). DNA was quantified using a QuBit^®^ 3.0 fluorometer (Invitrogen) with the dsDNA BR kit (broad range, 2 to 1000 ng/μL) according to the manufacturer’s protocol. The binding of large DNA fragments (117-8450 pb) was evaluated using gel electrophoresis (1.3% agarose), and the binding of small fragments (35-2000 bp) using a 2100 Bioanalyzer with an Agilent high-sensitivity DNA chip (Agilent Technologies).

**FIGURE 4:**
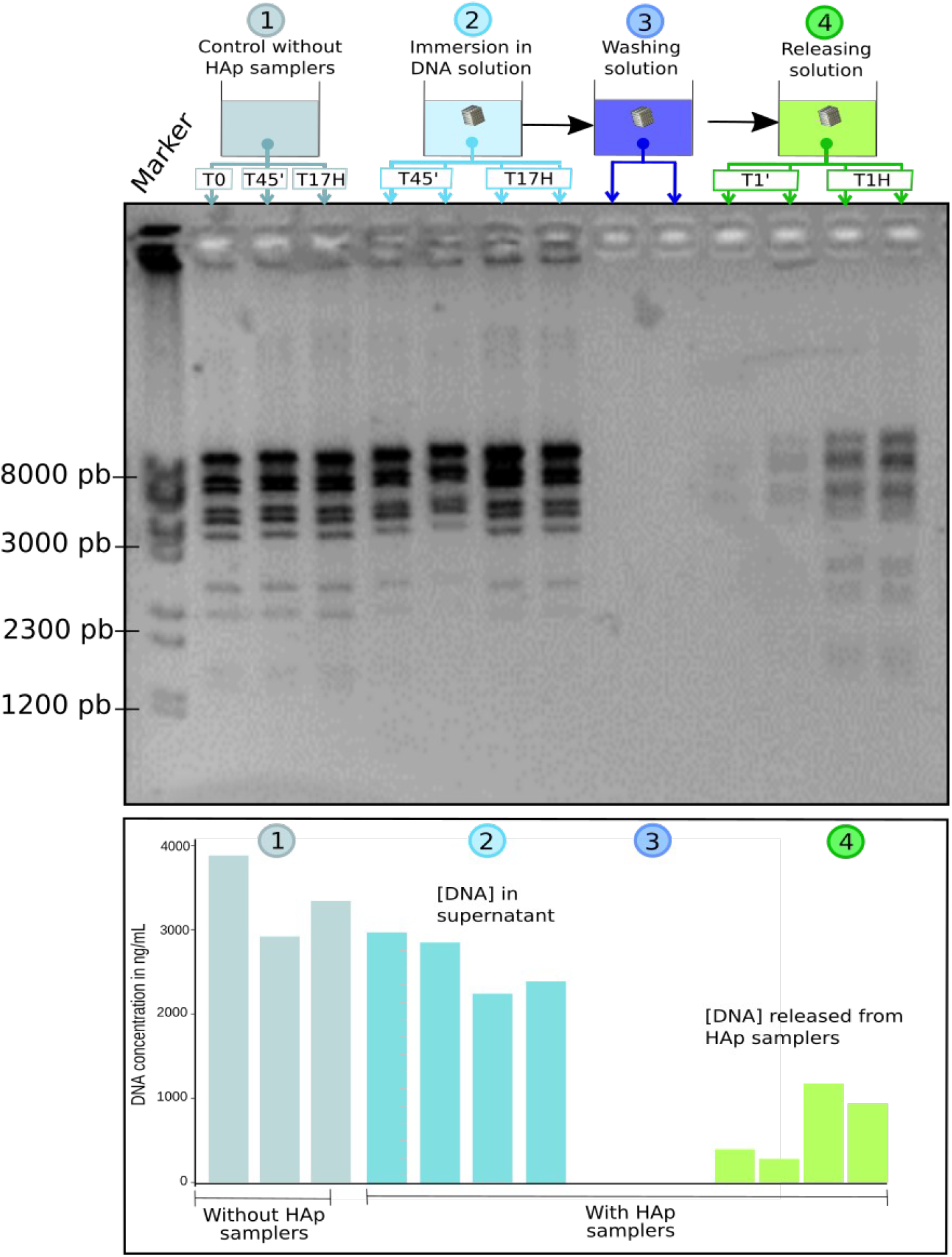
DNA binding and release by two replicates of the HAp samplers prototype 1. DNA fragment size (agarose electrophoresis gel, top panel) and concentration (bottom panel) are shown in the following order (left to right): 1) in the control solution without HAp samplers after 0 minute, 45 minutes and 17 hours, 2) in the DNA solution 45 min and 17 H after addition of HAp samplers, 3) in the washing solution and 4) in the releasing solution 1 min and 1 H after immersion of the HAp samplers.

For the second experiment (repeatability), the amount of DNA released from the HAp samplers was measured by fluorescence (excitation at 480 nm and emission at 520 nm) using an Infinite M200 Pro microplate fluorometer (TECAN, Switzerland). A QuantiFluor^®^ dsDNA kit was used according to the manufacturer’s protocol, with a DNA sample volume of 10 μL and 190 μL of working solution. A five-fold dilution series (1500-0 ng/μL) of standard DNA (Lambda DNA Standard, 100μg/mL) was used to build the standard curve and calculate the sample DNA concentration in μg/μL. The results are reported in percentage of recovered DNA (i.e. DNA bound and released).

*DNAr* is the measured concentration of released DNA (μg/mL) per HAp sampler and DNAtot is the initial DNA concentration added in each tube (DNAtot = 2.8 μg/mL).

#### 2.5.2 Quantitative PCR assay for eDNA from *A. aquaticus*

For the third experiment (microcosm experiment), quantitative PCR (qPCR) was used to quantify the amount of A.aquaticus eDNA recovered by the samplers. We designed a pair of primers to specifically amplify a 110 bp fragment of the mitochondrial 16S gene of *A. aquaticus (5’* GGTTTAAATGGCTGCAGTATCC 3’, 5’ CTTGTGTAATAAAAAGCCTACCTC 3’). The amplification specificity of the primers was tested *in silico* using primer-BLAST (NCBI) and assessed experimentally through PCR and electrophoresis gel analysis. The qPCR reaction volume was 20 μL consisting of 1X SsoAdvanced Universal SYBR Green Supermix (Bio-Rad Laboratories Inc., Hercules, CA), 0.5 μM of primers and 2 μL of DNA released from samplers. All qPCRs assays were run in duplicate in 96 wells plate on a CFX96 Touch™ RealTime PCR Detection System (Bio-Rad Laboratories, Inc., Hercules, CA). qPCR cycle started with an incubation at 95 °C for 10 min followed by 45 cycles of denaturation at 95 °C for 10 sec and an annealing/extension step at 60 °C for 20 sec before a final melt curve from 65-95 °C (0.5 °C increments). Each qPCR plate included a five-fold dilution series of the genomic DNA at a concentration between 0 and 2.5 ng/μL quantified by a QuBit 3.0 assay.

### 2.6 Statistical analysis

Linear mixed-effect models (LMMs) were used to test the influence of the prototype version and of the experiment timing (experiment 2). These models were fitted with the restricted maximum likelihood method using the package nlme. We tested significance of experiment timing and prototype version using likelihood ratio tests (LRT) between the models with and without the tested variable. All analyses were conducted using R (v 4.0.3).

## 3 Results

### 3.1 Experiment 1: DNA binding and release

We tested the hypothesis that 3D-printed HAp samplers can bind DNA of different fragment sizes by exposing them to two solutions of DNA size markers containing either long (117 to 8450 bp) or short DNA fragments (35 bp to 2000 bp). In the solution containing long DNA fragments, quantification of DNA concentration (Fig. 4, bottom panel) shows a clear reduction in DNA concentration in the supernatant after 17 hours of exposure to the HAp samplers for both sampler replicates. Once immersed in the releasing solution, and after only 1 minute, the HAp samplers started to release DNA. The amount of released DNA then tripled after 1 hour of incubation. By examining the DNA band profiles in the supernatants and in the releasing solution, we found that P1 bound all DNA fragment sizes from 2000 to 8450 bp. Fragments below 2000 bp were not visible on the electrophoresis gel (Fig. 4, top panel). The same observations were made on P2 (see supporting information).

Repeating the same experiment but using this time short DNA fragments (35-2000 pb) and a microfluidics-based automated electrophoresis system does not show an effect of fragment size on DNA binding (Fig. 5): both samplers prototypes bound and released DNA fragments ranging from 35 to 2000 bp although the resolution of the marker for fragments above 600 bp was not optimal.

**FIGURE 5:**
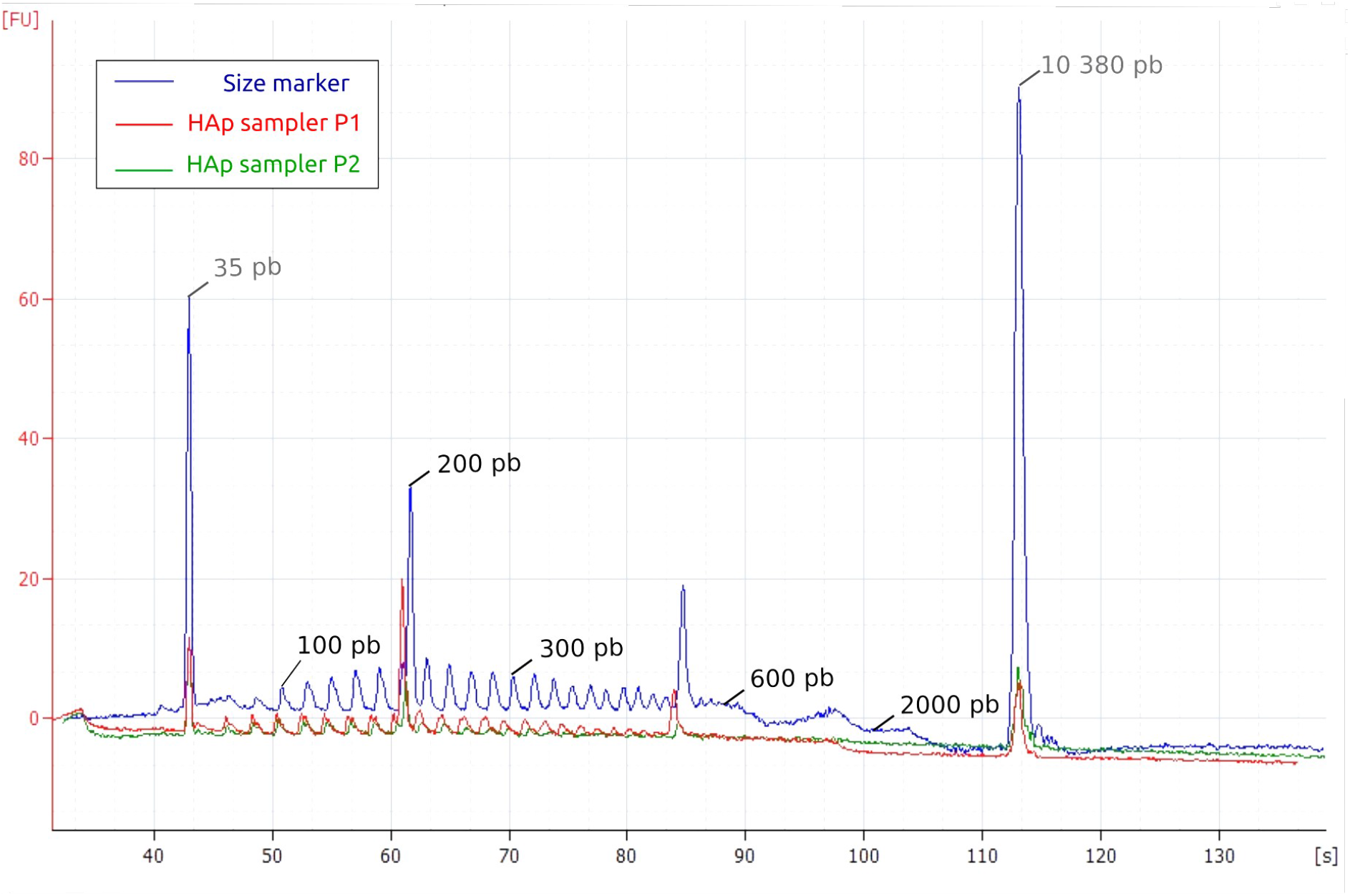
Electropherograms of the DNA fragments bound and released by the prototype 1 (red curve) and prototype 2 (green curve). As a standard, the blue curve represents the profile of the initial DNA marker. The horizontal axis represents the migration time of DNA fragments in seconds, and the vertical axis represents fluorescence. The left-most (35 bp) and rightmost (10380 bp) peaks are internal markers.

### 3.2 Experiment 2: Reusing HAp samplers over time

A repeatability experiment was conducted to test the hypothesis that HAp samplers can be reused and that their binding efficacy is stable after several cycles of use. We performed three consecutive cycles of use (experiment A, B and C), each composed of a decontamination, DNA binding and release steps. The percentage of DNA recovered by the samplers was lower in experiment A compared to experiments B and C, with an average of 8%, 17% and 15%, respectively (Fig. 6). In the meantime, experiment A showed a disproportion of samplers (18 out of 30, against 0 for experiment B and C) which failed to recover any DNA compared to the other experiments (Fisher exact test, p < 1E-10). After removing the samplers which failed to capture any DNA, we tested the influence of the experiment and prototype on the percentage of DNA recovered using a linear mixed-effect models with experiments (A, B and C) and sampler prototypes (P1 or P2) as the fixed effects, and samplers as random effect on the intercept. The experiment and the sampler prototype had no significant effect on the percentage of DNA recovered (LRT, sampler prototype: 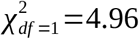, *p*=0.08, experiment: 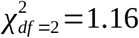, *p* = 0.28). Nonetheless, while not associated with any experiment in particular, the percentage of DNA recovered was highly variable. The coefficient of variation of DNA recovered was on average 65% considering all the samplers and 34% when excluding the samplers which failed to recover any DNA. Altogether, while we found that the samplers can still recover DNA after several cycles of use, we also discovered that the capacity of HAp samplers to recover DNA is variable and unpredictable: at some times it may not work at all, while at others it may recover a large amount of DNA.

**FIGURE 6:**
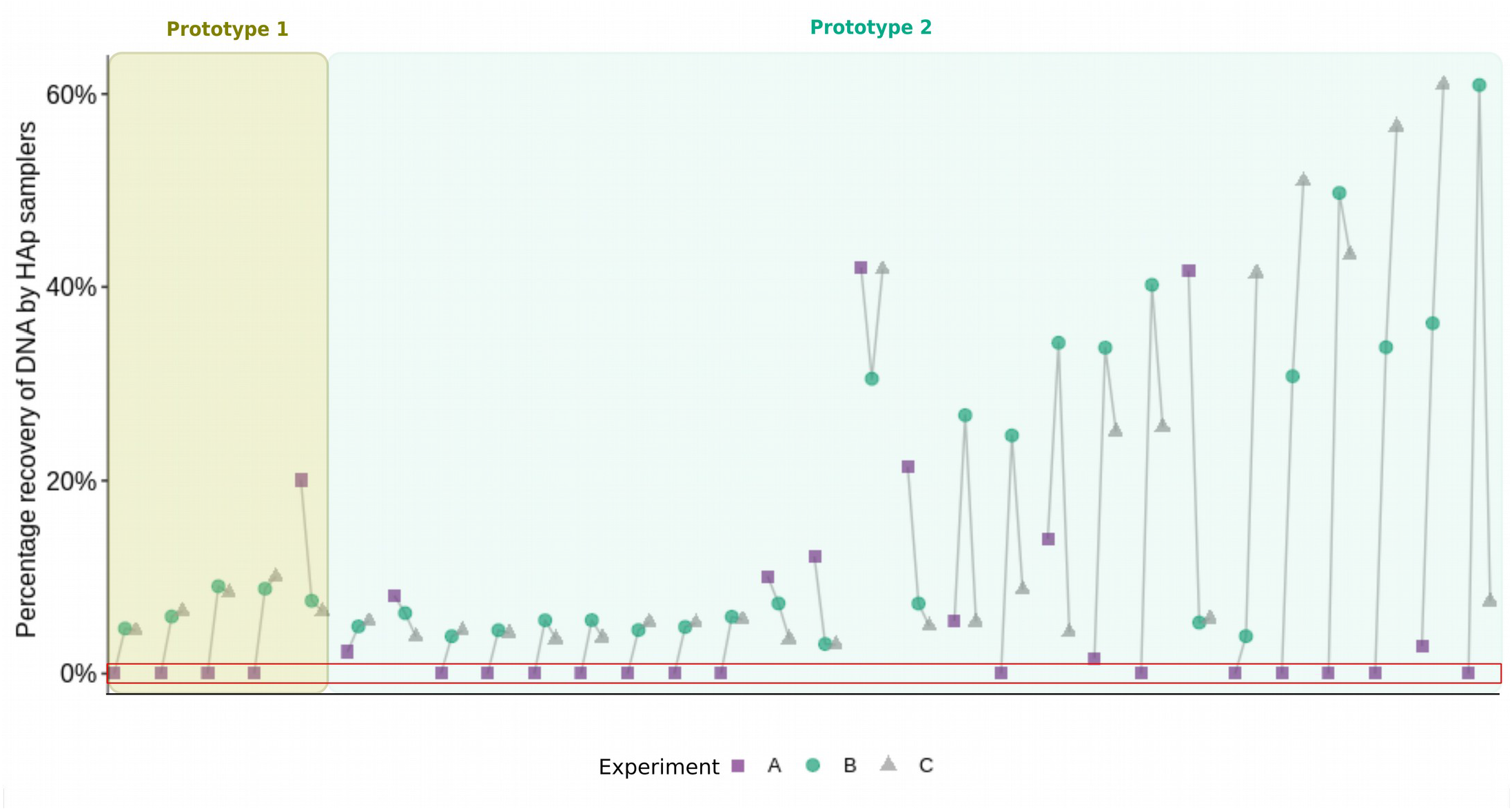
Percentage of DNA recovered by two prototypes (P1 and P2) of HAp samplers in three consecutive experiments (A, B and C). 5 P1 and 25 P2 samplers are sorted according to their variance of recovered DNA. Samplers in the red box did not recover any DNA.

### 3.3 Experiment 3: Environmental DNA sampling

We deployed the HAp samplers in a microcosm containing isopods (*Asellus aquaticus*) to test their ability to recover eDNA and used qPCR to quantify the amount of *A. aquaticus* 16S gene recovered by the samplers. In a microcosm with no organisms, we observed low levels of DNA that were similar or slightly above the amount of DNA observed in control samplers that were not immersed in a microcosm (Fig. 7). This is indicative of a slight level of crosscontamination between microcosms, and allowed us to determine an amount of 16S DNA below which we cannot differentiate between a contamination and a positive result (limit of blank, LOB). As expected, using concentrated genomic DNA as a positive control, the samplers recovered large amounts of 16S DNA molecules (Fig. 7). In the microcosm that contained isopods, the amount of 16S DNA molecules was about 3 orders of magnitude lower, with 3 samplers out of 8 below the limit of blank. Overall, the two HAp prototypes recovered *A. aquaticus* eDNA with the same efficiency (Wilcox-test, p = 0.89).

**FIGURE 7:**
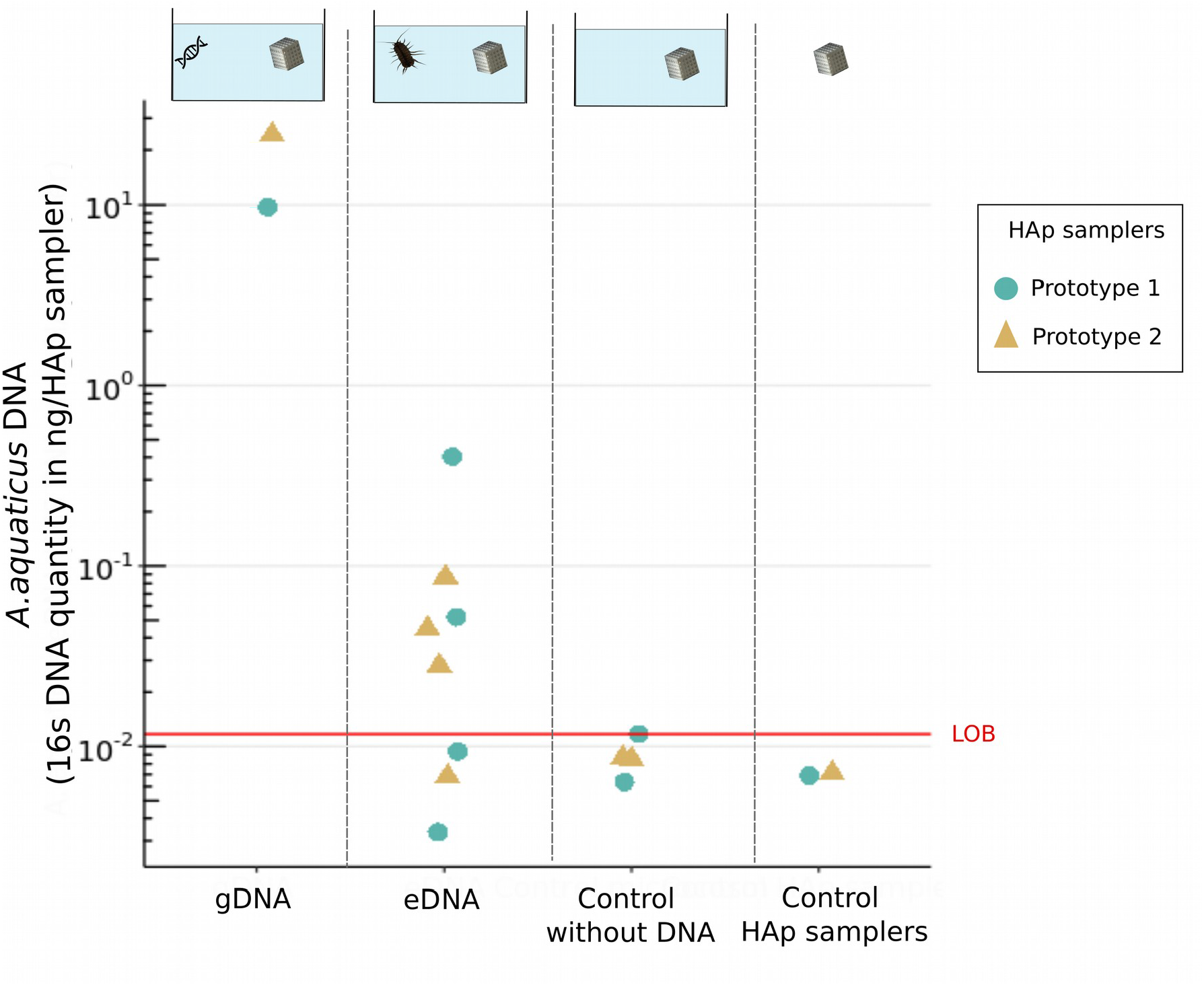
Quantity (in log scale) of *A. aquaticus* 16S gene recovered by the two prototypes of HAp samplers (P1 = circle, P2 = triangle) after 24h of incubation in microcosm containing genomic DNA used as a positive control (gDNA) or five individuals of *A. Aquaticus* (eDNA).

Two types of negative controls were used: one control microcosm without DNA (control without DNA), and HAp samplers without microcosm incubation (control HAp samplers). Red line corresponds to the limit of blanks (LOB).

## 4 Discussion

### 4.1 HAp samplers recover artificial DNA and eDNA

Our results demonstrated that HAp samplers can passively recover artificial DNA and eDNA. Using artificial DNA, DNA recovery was fast and optimal. In 17 hours, HAp samplers recovered up to 4 times more DNA (i.e. 1.75 μg) than the theoretical quantity we had estimated using a projection of a DNA monolayer on the surface of the samplers (see methods). These results confirm the high binding affinity between DNA and hydroxyapatite reported in literature (Okazaki et al., 2001; Del Valle et al., 2014) and suggest that more than one layer of DNA molecules can bind to the HAp surface. To our knowledge, this is the first time that these binding properties are tested and validated on 3D-printed objects. eDNA experiments showed that HAp samplers recovered eDNA from living macro-organisms (*A. aquaticus*). 5 out of 8 HAp samplers allowed a positive detection of *A. aquaticus* after only 24 hours of incubation in microcosm. Given the low densities of these small isopods which, unlike large organisms commonly used in eDNA microcosm experiments (e.g. fish, amphibians; Maruyama et al., 2014; Jo et al., 2020), are likely to release very small amounts of eDNA, and given the short experiment duration, this overall high rate of detection demonstrates the high sensitivity of HAp samplers to detect organisms.

### 4.2 Binding of different DNA fragment size

While it was hypothesized that DNA fragment size would influence DNA binding (Ogram et al., 1994), we did not find any evidence that certain fragment sizes bind preferentially to the HAp samplers. The samplers recovered DNA fragments of various sizes (i.e. 35-8450 bp), although bands below 2000 bp were not visible on the electrophoresis gel, possibly due to a higher concentration of the larger fragments in the marker solution. However, the sensitive microfluidics-based automated electrophoresis analysis showed that smaller fragments (<2000 bp) were bound and released by HAp samplers. eDNA is a complex mixture of genetic material ranging from cells to more or less degraded free DNA fragments (Wilcox et al., 2015). A sampling method that is not biased toward a given range of fragment sizes is a real advantage for eDNA sampling, in particular in environments where eDNA could be rapidly degraded into small free DNA fragments (Seymour et al., 2018). While free DNA binds to the HAp samplers, it remains to be tested whether other forms of eDNA such as proteo-nucleic complexes or even larger particles can also be collected.

### 4.3 Repeatability issues

Although HAp samplers show a great potential for DNA sampling, repeatability appears to be a concerning issue. Many HAp samplers showed extreme variability in DNA recovery among experiments carried out under strictly identical conditions (section 3.2). Given the high number of samplers that did not recover any DNA during the first but not the later experiments (Fig. 6), one might have expected that DNA recovery would improve with cycle of use. However, no effect of cycle of use or sampler prototype was found. In some cases, the DNA recovery remained stable over time, in some it increased, and in other it decreased. Surprisingly, while highly variable, there was not a set of samplers or one prototype in particular which was less effective than the others to recover DNA. This unexplained variability highlights the complexity of the binding mechanism between DNA and hydroxyapatite and the factor that controls it, and reinforces the necessity to better understand the evolution of the HAp surface after several DNA cycles of use. According to Okazaki et al. (2001), the binding affinity is based on an electrostatic interaction between the negative charges of the phosphate groups of DNA to the calcium ions at the surface of the hydroxyapatite. This ionic interaction strongly depends on the physico-chemical properties of the sampler surface and the solution in which the binding reaction takes place (Gallo et al., 2018). Among the surface properties, porosity, specific surface area, crystallinity and stoichiometry of the HAp phase (calcium groups can be substituted by other ions) could play a major role in DNA binding. The different manufacturing steps, such as the HAp densification (i.e. sintering), can greatly influence most of these surface properties. In particular, ionic substitution (e.g. carbonatation) and partial dehydration are known to occur frequently in HAp during thermal treatment (Wang, Dorner-Reisel and Müller, 2004; Lafon, 2004) such as the ones used here to decontaminate the samplers before and between experiments, and might be the source of the observed variability. Surface analyse needs to be carried out to identify the physical (e.g. porosity, crystalline phases) and chemical (e.g. surface ionic groups) parameters involved in DNA binding on the HAp surface and the extent to which these parameters are influenced by the manufacturing and use of the sampler (e.g. sintering, debinding, immersion in DNA solution).

## 5 CONCLUSION

In view of the democratisation of the use of eDNA, tools are needed to easily and cost-effectively sample eDNA. We demonstrate that 3D passive hydroxyapatite samplers can be designed and used to collect eDNA, albeit some repeatability issues. Provided we can get a better understanding and control of the interaction between eDNA and HAp, this approach offers an alternative sampling solution for eDNA-based biomonitoring. It also opens up an interdisciplinary field at the interface between engineering, surface science and molecular ecology.

## ACKNOWLEDGEMENTS

This work was supported by the CNRS Mission pour les Initiatives Transverses et Interdisciplinaires (project XLIFE CAPTAS), the French Biodiversity Agency (OFB), the National Technology Research Association (ANRT) and the company Eurofins Hydrobiologie France. This work was realised thanks to the support of the Graduate School H_2_O’Lyon (ANR-17-EURE-0018) and Université de Lyon (UdL) as part of the programme “Investissements d’Avenir” run by Agence Nationale de la Recherche (ANR). We acknowledge Louise Camus for her help with the microcosm experiment, Valentin Vasselon for his advice on experiments with artificial DNA and Jalal Omarakly for the surface analysis of the samplers.

## AUTHORS’ CONTRIBUTIONS

TL and CM conceived the ideas and designed HAp samplers. Experimental design was conceived by TL, LK and HV. HAp samples were thermal treated and characterized by HR, ST and LG. Laboratory experiments were conducted by HV and LK. Data analysis was conducted by HV and TL. HV and TL led the writing of the manuscript. All authors contributed to the manuscript.

## Notes

### Competing Interest Statement

The authors have declared no competing interest.

