## Supplemental_Figure_A for "Passive sampling of environmental DNA in aquatic environments using 3D-printed hydroxyapatite samplers"

**SUPPORTING INFORMATION**

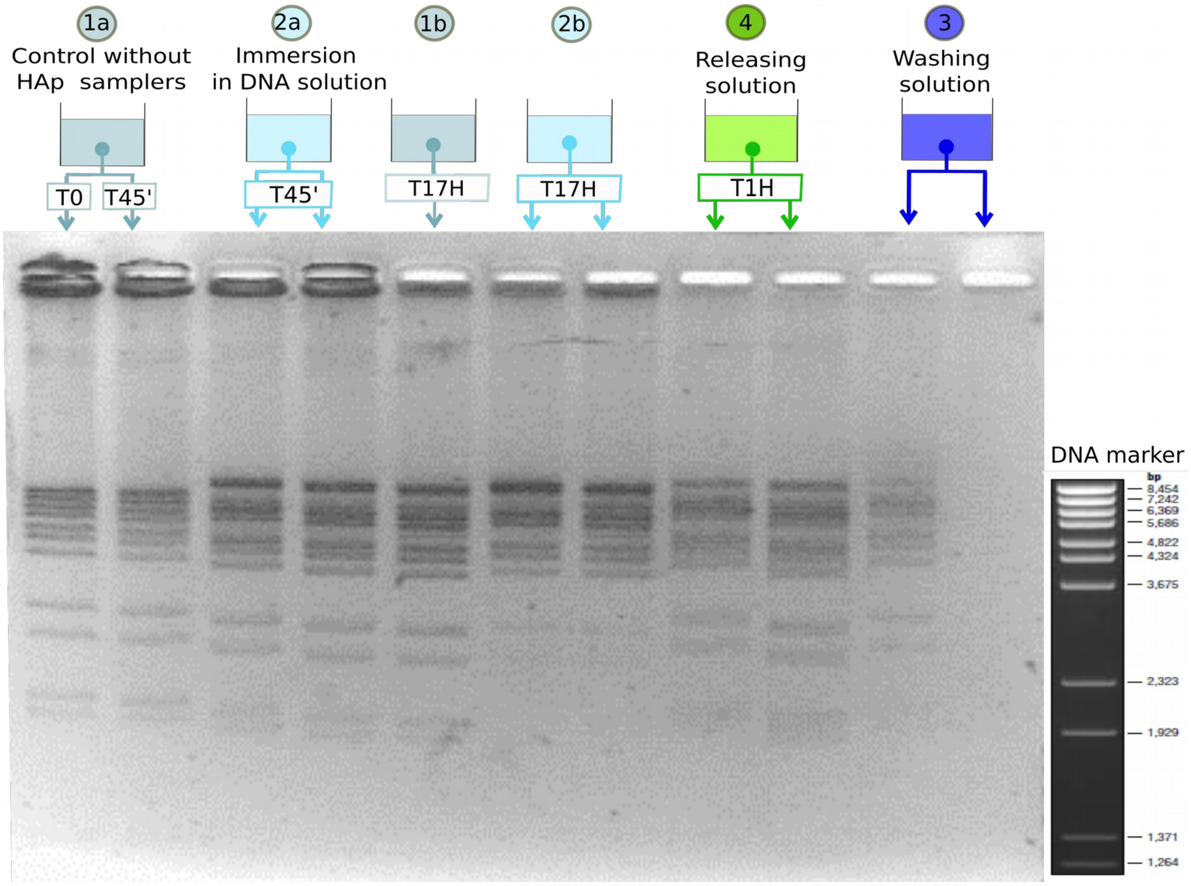


**FIGURE A** : DNA binding and release by two replicates of the HAp samplers prototype 2. DNA fragment size is shown in the following order : 1a) in the control solution without HAp samplers after 0 minute and 45 minutes, 2a) in the DNA solution 45 min after addition of HAp samplers, 1b) in the control solution without HAp samplers after 17H, 2b) in the DNA solution 17H after addition of HAp samplers, 3) in the releasing solution 1 H after immersion of the HAp samplers and 4) in the washing solution. DNA was stained directly in the samples with GelRed. As a result, the same DNA fragment in two samples but at two different concentrations will migrate at different speed.
